# Early dosage compensation of zygotically-expressed genes in *Drosophila melanogaster* is mediated through a post-transcriptional regulatory mechanism

**DOI:** 10.1101/2022.04.25.489409

**Authors:** Victoria M. Blake, Michael B. Eisen

## Abstract

Many key regulators of early embryogenesis in *Drosophila melanogaster* are X-linked. However, the canonical, MSL-mediated dosage compensation, which involves hyper-transcription of the genes on the single X chromosome in males, is not active until the post-syncytial stage of development. A separate MSL-independent dosage compensation system active earlier in development has been described, though the mechanism through which the process functions remain unclear. In this study, we quantified transcription in living embryos at single-locus resolution to determine if early dosage compensation of the X-linked genes *buttonhead* and *giant* is sensitive to X chromosome dose. We found no evidence for a transcriptionally regulated mechanism of early dosage compensation, suggesting that the previously observed compensation of mRNA levels for these genes is achieved via a post-transcriptional regulatory mechanism.

## Introduction

In multicellular diploid organisms with chromosomal sex determination, the functional degeneration of sex chromosome specific to the heterogametic sex (the Y in XY systems and W in ZW systems) creates a dosage imbalance for sex-chromosome encoded genes. To mitigate the detrimental impact that this dosage imbalance could have on fitness, a variety of mechanisms have evolved to equalize the abundance of gene products found on only one sex chromosome (typically the X or Z).

In the vinegar fly *Drosophila melanogaster*, which uses an XX/XY sex-determination system, dosage compensation is achieved by upregulating transcription of genes on the male X chromosome (reviewed in Kuroda et al. 2016). Mutations that affect this process are lethal in males (Belote and Lucchesi 1980, Lucchesi and Skripsky 1981, Hilfiker et al. 1997, Meller and Ratner 2002).

All but one of the Male Sex Lethal (MSL) proteins that mediate this process are present in the maternally deposited protein component. Male specificity is achieved through the activity of the male-specific MSL-2, which directs assembly of the dosage compensation complex to the X chromosome, resulting in acetylation of H4K16 by MOF (Hilfiker et al. 1997, Smith et al. 2000) and a subsequent doubling of transcription from X-linked genes (Smith et al. 2001, Gelbart et al. 2009).

However, an early, MSL-2 independent dosage compensation system acting on *runt* was discovered via genetic experiments (Gergen 1987, Bernstein and Cline 1994) and shown to apply more broadly by direct measurement of mRNA levels in early male and female embryos (Lott et al. 2011). It was hypothesized, based on the enrichment of putative binding sites for the master sex regulator Sxl in the 5’ and 3’ untranslated regions of X encoded genes, that this system was based on post-transcriptional regulation of mRNA levels (Bernstein & Cline 1994, Lott et al. 2011).

Here we use quantitative live-embryo imaging (Garcia et al. 2013, Bothma et al. 2014) to compare rates of transcription at individual X-linked loci across early development in male and female embryos. Briefly, transcription of a gene tagged with a reporter sequence (MS2 or PP7) forms stem loops that, when bound by a coat protein (MCP or PCP) conjugated to GFP, gives rise to a fluorescent signal. This allows for simultaneous direct quantification via microscopy of transcription rates at individual loci across many nuclei. The ability to quantify transcription rates of candidate genes in embryos of both sexes makes this system uniquely amenable for testing if early dosage compensation is due to increasing the rate of transcription of genes on the male X chromosome.

We focus on four X-linked genes: *giant* (*gt*) and *buttonhead* (*btd*) that had previously been shown to be dosage compensated in the early embryo, and *embryonic lethal abnormal vision* (*elav*) and *bangles and beads* (*bnb*) whose early mRNA levels are dosage sensitive. We found that the early dosage compensation observed for these genes was not due to a doubling of transcription in males, suggesting that this process is likely regulated post-transcriptionally. In addition, we discovered an unreported doubling of the transcription rate in females for *elav*, a gene that is not dosage compensated and has no defined role in sex differentiation.

## Materials and Methods

### DNA constructs

Fly strains were tagged with MS2 or PP7 reporter sequences at the endogenous loci of X-linked genes through CRISPR-Cas9 Homology-Directed Recombination. We used the U6-gRNA protocol from flycrispr.org (Gratz et al. 2016) to clone three guide RNA sequences per gene, targeting a 100 bp region around the start codon, into pCFD5 plasmids (Port and Bullock 2016). For donor plasmids, we inserted a 24X MS2 or PP7 cassette (Garcia et al. 2013, Liu et al. 2020) between 1kb of homology arms for a gene of interest into a pUC19 vector backbone using the NEBuilder HiFi DNA Assembly protocol. We attempted to tag all target genes with both MS2 and PP7, but, due to low editing efficiency and potentially other factors, recovered only one for each gene, as described in text.

### Fly line generation

gRNA, donor, and selection plasmids (for phenotypic screening, Kane et al. 2017) were purified and sent to Rainbow Transgenic Inc. for microinjection into *y2 cho2 v1; Sp/CyO, P{nos-Cas9, y+, v+}2A* embryos.

Full details of construct and sequence information can be found in a public <u>Benchling folder</u>.

Progeny of injected flies with an *ebony* phenotype were molecularly screened by PCR for the insertion of the MS2 or PP7 cassettes at endogenous X-linked loci. Lines with successful transgene insertion were confirmed by Sanger sequencing.

Transcription of X-linked genes was measured by imaging embryos homozygous for the MS2- or PP7-tagged gene and a constitutively expressed *Histone-RFP; MCP-eGFP* (Garcia et al. 2013) or *PCP-eGFP* (Larson et al. 2011) allele.

### Live imaging

#### Image acquisition

Sample preparation followed the procedures described in (Bothma et al. 2014, Garcia et al. 2013, Lammers et al. 2020). In brief, embryos were collected, dechorinated with 50% bleach, and mounted between a semipermeable membrane (Lumox film, Starstedt, Germany) and a coverslip while in Halocarbon 27 oil (Sigma). Data collection was performed using a Zeiss LSM 800 scanning confocal microscope (Zeiss, Jana, Germany). The MCP-eGFP or PCP-eGFP were excited with laser wavelengths of 488 nm, respectively. Data were collected at a 40X objective with oil immersion where the average laser power on the specimen was 7.3 mW and a master gain of 550V. The confocal stack consisted of 21 equidistant slices with an overall z-height of 5.97μm and an interslice distance of 0.29 μm. The images were acquired at a frame time of 633.02 ms and a pixel dwell time of 1.03 μs. Image sizes were 78 μm x 19.5 μm, a frame size of 1024 pixels x 256 pixels, and a pixel size of 0.08 μm. Data were taken for at least 3 embryos of each sex per genotype, and each embryo was imaged for at least the first 30 minutes of nuclear cycle 14.

#### Image analysis

Live-imaging data were analyzed using a custom-written software following the protocols in (Garcia et al. 2013, Lammers et al. 2020). This software, containing MATLAB code, can be found in a <u>public GitHub repository</u>. In brief, this procedure involves nuclear segmentation, segmenting transcription spots based on fluorescence, and calculating the intensity of each MCP-eGFP or PCP-eGFP transcription spot inside a nucleus as a function of time.

#### Data analysis

To infer bursting parameters from experimental fluorescence traces, we used a compound-state hidden Markov Model described in Lammers et al. 2020 whose code can be found in a <u>public GitHub repository</u>.

All data analysis was done in Python using a Jupyter Notebook with custom code to generate figures. The Jupyter notebook and all data required to run it are available in Supplementary File 1 this <u>Github link</u>.

### Quantitative RNA-FISH

#### Probe design and hybridization

Custom Stellaris® FISH Probes were designed against *gt* and *bnb* RNA by utilizing the Stellaris® RNA FISH Probe Designer (Biosearch Technologies, Inc., Petaluma, CA) available online at www.biosearchtech.com/stellarisdesigner (version 4.2). Single-molecule RNA-FISH protocol was followed using the *D. melanogaster* embryo protocol at the Biosearch Technologies website.

#### Image acquisition

Embryos were staged at nuclear cycle 14A based on percent membrane invagination during cellularization (< 25%). Data collection was performed using a Zeiss LSM 800 scanning confocal microscope (Zeiss, Jana, Germany). A laser wavelength of 670 nm was used to excite the probe-conjugated fluors. Data were collected at a 63X objective with oil immersion with the laser power on the specimen was set to 5% and a master gain of 650V. The images were acquired at a frame time of 930.91 ms and a pixel dwell time of 1.52 μs. Image sizes were 202.8 μm x 202.8 μm, a frame size of 512 pixels x 512 pixels, and a pixel size of 0.4 μm.

The field of view was adjusted to encompass the entire expression domain of probed patterned genes, and the bounds of the z-stack for each image were determined by when the z-plane no longer detected the probe fluorescent signal.

Full details of probe sequence information can be found in a public <u>Benchling folder</u>. Raw and max intensity projection images can be found in Supplemental File 2.

#### Image analysis

Total signal of mRNA with background correction was measured for probed genes using ImageJ. Z-slices of the original image that contained no signal were trimmed from the image before generating a 2D max intensity projection. We followed the Cooper Lab protocol for Quantitation of Total Fluorescence per Cell with Background correction (https://cooperlab.wustl.edu/). In brief, two ROIs were determined in ImageJ: one to measure signal and the other to measure background. We then averaged the mean Integrated Density of the signal (5 separate measurements encompassing the total expression domain) and of the background. The mean background is subtracted from the mean integrated density, and then divided by the area of the signal ROI to obtain the Corrected Integrated Density.

## Results

Previous work identified a wide-spread dosage compensation of zygotic transcripts prior to cellularization, where 36 out of 85 X-linked genes showed less than a 1.5-fold excess of transcript abundance in female embryos relative to their male counterparts (Lott *et al.,* 2011). Many of these early-compensated genes are key developmental regulators, where differences in transcript abundance could pose profound consequences to organismal development. Because these early-compensated genes are regulated before MSL-2, the molecule that mediates canonical dosage compensation, is expressed, we sought to investigate if the early-compensation of these developmental regulators occurred through a transcriptional mechanism independent of MSL-2.

We chose two early-compensated genes (*gt* and *btd*) for further study that are essential for mediating embryonic development and whose mRNA abundance is matched in male and female embryos during the early nuclear cycles. Because not all zygotically-transcribed genes emanating from X are early-compensated, we chose two additional genes (*elav* and *bnb*) with twice as much female mRNA abundant relative to males as a point of comparison in our study. All four of these genes reach maximum transcript abundance during the first half of nuclear cycle 14 (Lott *et al.,* 2011), mitigating any differences due to developmental stages across datasets.

We used CRISPR-Cas9 (Gratz *et al.,* 2015) to insert an array of either MS2 or PP7 stem loop sequences immediately upstream of the translation start codon of the endogenous gene of interest. The rationale behind choosing this insertion site was to position at the stem loops sequences at beginning of the resulting transcript, ensuring a robust reporter signal of when transcription is occurring. The MS2 or PP7 sequences were flanked by the donor and acceptor splice sites from the *hunchback* intron so that the stem loops would be removed from the mature mRNA to prevent problems with RNA localization (Heinrich *et al.,* 2017) or the translation of the endogenous protein.

We homozygosed the lines containing MS2 and PP7-tagged genes and crossed in a transgene encoding an MCP-GFP (or PCP-GFP) fusion protein that binds transcribed stem loop sequences. The binding of fluorescently conjugated coat protein to these stem loops (Bernardi *et al.,*1972) gives rise to a fluorescent puncta (what we will refer to as “spots” moving forward) in the nucleus (Figure 1B; Garcia *et al.,* 2013) whose fluorescence intensity correlates with the abundance of RNA present at the locus (Garcia et al. 2013). This system offers a spatio-temporal and quantitative characterization of transcript abundance for a gene of interest (Movies 1 & 2).

**Figure 1:**
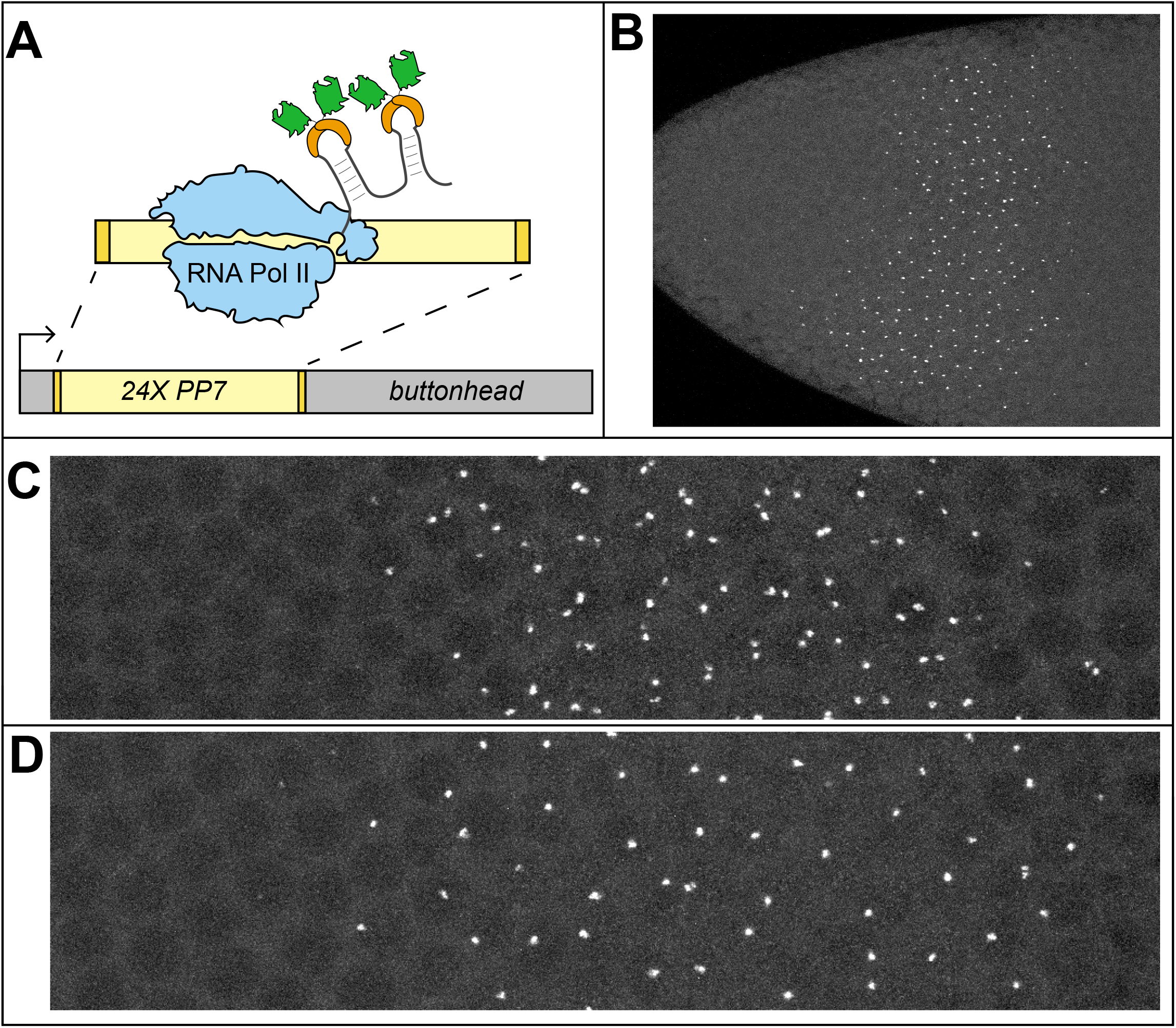
MS2-MCP and PP7-PCP reporter systems measure transcription rate of X-linked genes in the early *Drosophila melanogaster* embryo. A) An example schematic of how the PP7-PCP reporter system works: 24 repeats of the PP7 stem loop sequence (yellow) are flanked by splice donor and acceptor sites from the intron of the *hunchback* gene (gold) were inserted upstream of the target gene’s start codon (ATG), represented here by the gene “*buttonhead”*. When transcribed, the PP7 form RNA step loops that are subsequently bound by PP7 Coat Protein (orange) conjugated to GFP (green). B) The anterior portion of an embryo expression *PP7-buttonhead* (*PP7-btd*). C) Zoomed-in view of the *PP7-btd* expression domain in a female embryo, made evident by the two transcribing loci in nuclei as opposed to D) the same field of view for *PP7-btd* expression in a male embryo, where there is only one transcribing locus per nucleus.

Embryos with successfully incorporated MS2-MCP or PCP-PP7 reporter systems showed no signs of impaired organismal health and faithfully recapitulated the tagged gene’s expression patterns previously reported by FISH (Figure 1B; Wimmer *et al.,* 1996, Tomancak et al. 2002). A key feature of this system that lends itself for studying dosage compensation is the ease in sexing embryos–the fluorescent spots in nuclei serve as a readout for the number of X chromosomes present (embryos with two fluorescent spots within the same nucleus are female, while embryos with one spot are male) (Figure 1C & 1D).

We used laser-scanning confocal microscopy to acquire MCP-GFP (or PCP-GFP) movies of embryos during nc14, the last nuclear cycle before gastrulation. To ensure consistency across fields of view across these high magnification movies, we optimized data collection for patterned genes to capture all nuclei along the AP axis within the same single expression domain (Movies 1 & 2, Supplemental Figure 1). If a gene had a more ubiquitous expression domain within the anterior domain of the embryo, we positioned the anterior pole relative to the field of view consistently across all data collections. To obtain high temporal resolution and optimize signal-to-noise with minimal bleaching, we collected movies so that the acquisition time for each timepoint (or Z-stack) corresponded to intervals of 19.5s. In total, we collected 51 movies (Movies 3-51) with a minimum of three movies of each sex per genotype (see Table 1).

**Table 1:** MS2 or PP7 live-imaging data collected in this study

| Embryo ID | Gene | Sex | nuclear cycle | Duration (in frames) | Total time | Movie # |
| --- | --- | --- | --- | --- | --- | --- |
| M_btd_1 | PP7-btd | M | 14 | 159 | 51.675 | Movie 1 |
| F_btd_1 | PP7-btd | F | 14 | 172 | 55.9 | Movie 2 |
| M_btd_2 | PP7-btd | M | 14 | 142 | 46.15 | Movie 3 |
| M_btd_3 | PP7-btd | M | 14 | 162 | 52.65 | Movie 4 |
| M_btd_4 | PP7-btd | M | 14 | 125 | 40.625 | Movie 5 |
| M_btd_5 | PP7-btd | M | 14 | 141 | 45.825 | Movie 6 |
| M_btd_6 | PP7-btd | M | 14 | 183 | 59.475 | Movie 7 |
| M_btd_7 | PP7-btd | M | 14 | 99 | 32.175 | Movie 8 |
| M_btd_8 | PP7-btd | M | 14 | 110 | 35.75 | Movie 9 |
| M_btd_9 | PP7-btd | M | 14 | 94 | 30.55 | Movie 10 |
| F_btd_2 | PP7-btd | F | 14 | 122 | 39.65 | Movie 11 |
| F_btd_3 | PP7-btd | F | 14 | 194 | 63.05 | Movie 12 |
| F_btd_4 | PP7-btd | F | 14 | 208 | 67.6 | Movie 13 |
| F_btd_5 | PP7-btd | F | 14 | 123 | 39.975 | Movie 14 |
| F_btd_6 | PP7-btd | F | 14 | 176 | 57.2 | Movie 15 |
| F_btd_7 | PP7-btd | F | 14 | 166 | 53.95 | Movie 16 |
| F_btd_8 | PP7-btd | F | 14 | 173 | 56.225 | Movie 17 |
| F_btd_9 | PP7-btd | F | 14 | 119 | 38.675 | Movie 18 |
| M_gt_1 | MS2-gt | M | 14 | 119 | 38.675 | Movie 19 |
| M_gt_2 | MS2-gt | M | 14 | 148 | 48.1 | Movie 20 |
| M_gt_3 | MS2-gt | M | 14 | 125 | 40.625 | Movie 21 |
| M_gt_4 | MS2-gt | M | 14 | 114 | 37.05 | Movie 22 |
| F_gt_1 | MS2-gt | F | 14 | 111 | 36.075 | Movie 23 |
| F_gt_2 | MS2-gt | F | 14 | 143 | 46.475 | Movie 24 |
| F_gt_3 | MS2-gt | F | 14 | 147 | 47.775 | Movie 25 |
| F_gt_4 | MS2-gt | F | 14 | 139 | 45.175 | Movie 26 |
| M_bnb_1 | PP7-bnb | M | 14 | 102 | 33.15 | Movie 27 |
| M_bnb_2 | PP7-bnb | M | 14 | 122 | 39.65 | Movie 28 |
| M_bnb_3 | PP7-bnb | M | 14 | 122 | 39.65 | Movie 29 |
| F_bnb_1 | PP7-bnb | F | 14 | 160 | 52 | Movie 30 |
| F_bnb_2 | PP7-bnb | F | 14 | 146 | 47.45 | Movie 31 |
| F_bnb_3 | PP7-bnb | F | 14 | 157 | 51.025 | Movie 32 |
| M_elav_1 | MS2-elav | M | 14 | 133 | 43.225 | Movie 33 |
| M_elav_2 | MS2-elav | M | 14 | 148 | 48.1 | Movie 34 |
| M_elav_3 | MS2-elav | M | 14 | 128 | 41.6 | Movie 35 |
| F_elav_1 | MS2-elav | F | 14 | 139 | 45.175 | Movie 36 |
| F_elav_2 | MS2-elav | F | 14 | 159 | 51.675 | Movie 37 |
| F_elav_3 | MS2-elav | F | 14 | 141 | 45.825 | Movie 38 |
| M13_btd_1 | PP7-btd | M | 13 | 33 | 10.725 | Movie 39 |
| M13_btd_2 | PP7-btd | M | 13 | 31 | 10.075 | Movie 40 |
| M13_btd_3 | PP7-btd | M | 13 | 25 | 8.125 | Movie 41 |
| F13_btd_1 | PP7-btd | F | 13 | 26 | 8.45 | Movie 42 |
| F13_btd_2 | PP7-btd | F | 13 | 23 | 7.475 | Movie 43 |
| F13_btd_3 | PP7-btd | F | 13 | 23 | 7.475 | Movie 44 |
| M13_bnb_1 | PP7-bnb | M | 13 | 38 | 12.35 | Movie 45 |
| M13_bnb_2 | PP7-bnb | M | 13 | 25 | 8.125 | Movie 46 |
| M13_bnb_3 | PP7-bnb | M | 13 | 29 | 9.425 | Movie 47 |
| F13_bnb_1 | PP7-bnb | F | 13 | 50 | 16.25 | Movie 48 |
| F13_bnb_2 | PP7-bnb | F | 13 | 44 | 14.3 | Movie 49 |
| F13_bnb_3 | PP7-bnb | F | 13 | 32 | 10.4 | Movie 50 |

We used a custom image analysis pipeline (Garcia et al. 2013, Lammers et al. 2020) to identify nuclei and extract fluorescent intensity measurements for actively transcribing X-linked loci in embryos of both sexes. A representative example of the resulting data is shown in Figure 2.

**Figure 2:**
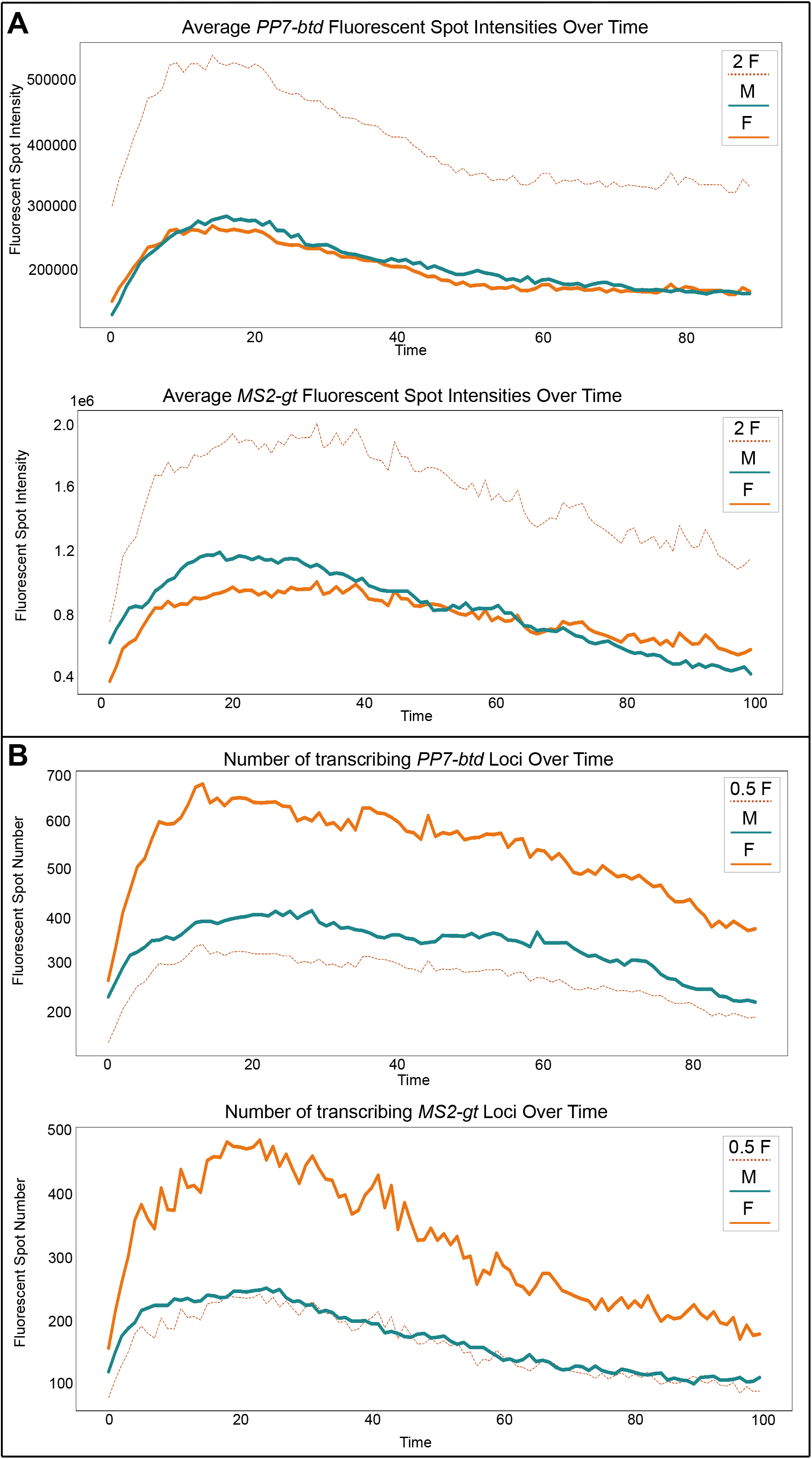
Transcription rate of the early dosage compensated *PP7-btd* and *MS2-gt* genes is the same in male and female embryos. A) Average distributions of fluorescent spot intensity over time (in frames, where “1” is the first instance of transcription of the locus during nuclear cycle 14) for all female (orange) and male (dark cyan) *PP7-btd* (N=9 embryos of each sex) and *MS2-gt* (N=4 of each sex) embryos. The dashed, dark orange line represents double the average female fluorescent spot intensity for every given timepoint. B) The distribution of fluorescent spot intensities plotted over time for the dosage compensated *MS2-gt* (N=4 embryos of each sex) with total spot intensity histograms on the right. B) The number of transcribing *PP7-btd* and *MS2-gt* loci throughout nuclear cycle 14. The dashed orange line represents half of the transcribing female loci.

### No hyper-transcription of male X in dosage compensated genes

We compared the average fluorescence intensity (which we expect to be proportional to polymerase density on transcribing loci) of detected transcription foci from male and female X chromosomes across time during mitotic cycle 14 for both *btd* and *gt* (Figure 2A). For *btd* they are nearly identical, and for *gt*, the differences both balances out over time (with slightly higher transcription from male X’s early in cycle 14 and slightly higher transcription from female X’s later in cycle 14). As it is possible to increase transcriptional output even with identical polymerase density on transcribing loci by increasing the fraction of loci actively transcribing, we also looked at the number of transcribing loci as a function of time in male and female embryos (Figure 2B), and, as expected, observed them to be in approximately a 2:1 ratio between females and males. These results are inconsistent with early dosage compensation of *btd* and *gt* being achieved via hyper-transcription of the male X.

We made similar comparisons between average fluorescence and number of transcribing loci in the two non-compensated loci bnb and elav. These show more variation between sexes than two dosage compensated genes, with bnb generally higher in males, and elav higher in females (Figure 3).

**Figure 3:**
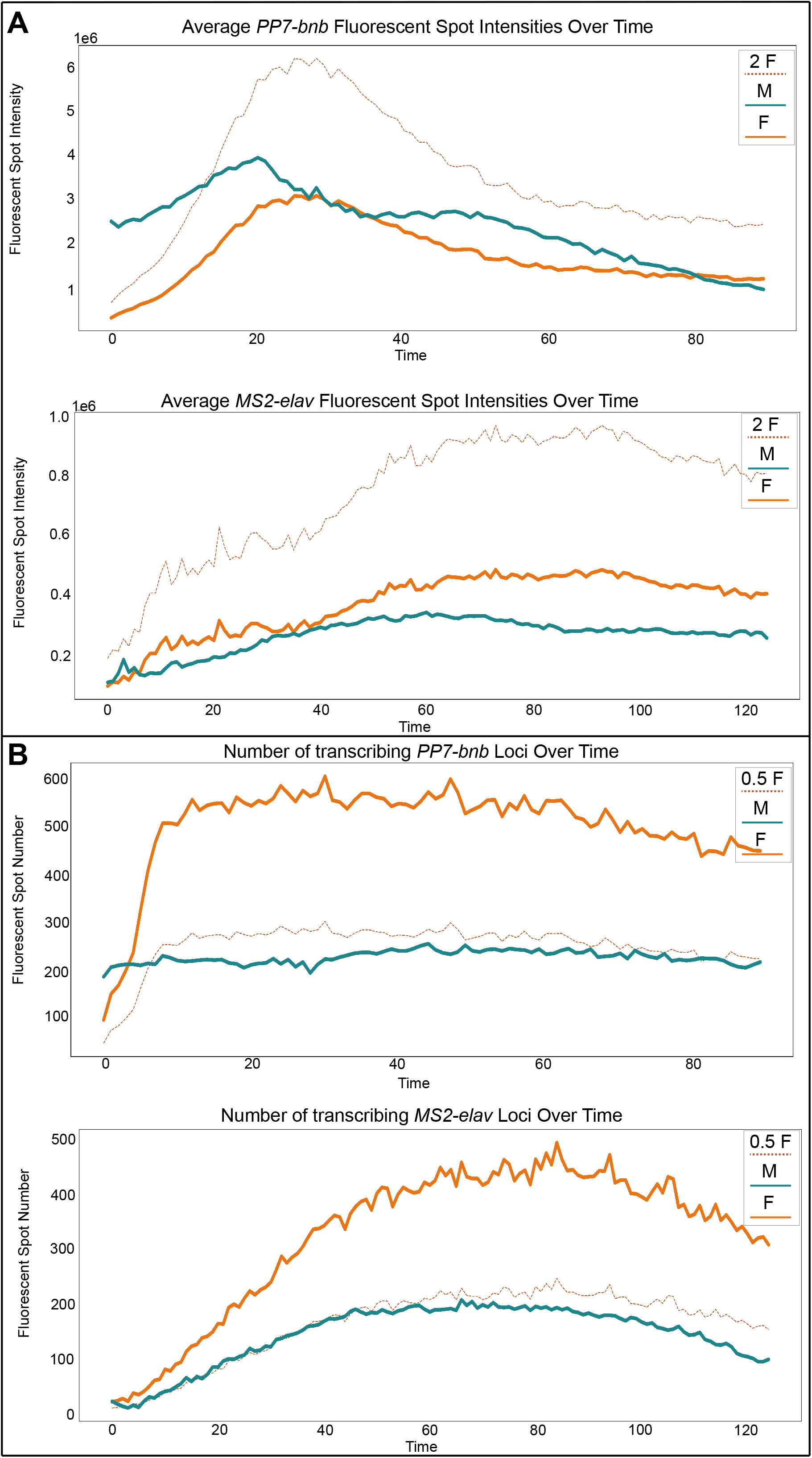
Transcription rate of *PP7-bnb* and *MS2-elav* is does not result in dosage compensation. A) Averages for male (dark cyan) and female (orange) *PP7-bnb* and *MS2-elav* fluorescent spot intensities plotted over nuclear cycle 14. The dashed, dark orange line represents double the average female fluorescent spot intensity for every given timepoint. B) The number of transcribing *PP7-bnb* and *MS2-elav* loci throughout nuclear cycle 14. The dashed orange line represents half of the transcribing female loci.

## Discussion

Our tailoring of quantitative live-imaging methods to examine early embryonic dosage compensation in *Drosophila melanogaster* were motivated by the hypothesis that the equalizing of mRNA abundance between sexes in the absence of MSL-2 mediated by regulation of transcription rate at X-linked loci.

Instead, what our studies revealed was a post-transcriptional mechanism for regulating early dosage compensation of two key developmental regulators, *btd* and *gt.* These findings represent a broadening of our understanding of different ways in which the sex chromosome dosage imbalance is mitigated during early embryogenesis. Our findings offer a more dynamic way of understanding how mRNA abundance is regulated across developmental stages. Importantly, this presents a shift in our understanding of how mRNA abundance is regulated considering the decades of research whose findings posited that dosage compensation is almost exclusively a transcriptionally regulated process.

Sex lethal (Sxl), and RNA-binding protein that inhibits dosage compensation in females, may play a role in regulating the abundance of X-linked mRNAs during early embryogenesis. The regulation of X-linked *runt* abundance by Sxl is the only documented observation of an MSL-2-independent, post-transcriptional regulation of dosage compensation prior to our study (Gergen 1987, Bernstein and Cline 1994). It is possible that because the two molecules regulate each other’s expression (Kramer et al. 1999, Torres et al. 2009), that observation of post-transcriptional dosage compensation by Sxl is gene-specific and therefore inapplicable to the widespread phenomenon we observe during early embryogenesis (Lott et al. 2011).

Considering our findings, the observed enrichment of predicted Sxl binding sites (poly-U tracts) in the 5’ and 3’ UTRs of many X-linked genes (Kelley et al. 1995) may point to one mechanism for post-transcriptionally regulating dosage compensation. There is no direct correlation between genes that are early dosage compensated and their predicted regulation by Sxl–not all genes that are early dosage compensated are predicted to be Sxl targets, and not all predicted Sxl targets are early dosage compensation. There may however be a subset of early dosage-compensated genes whose abundance may be regulated by Sxl, a hypothesis worth exploring in future studies.

It is formally possible that regulation of transcriptional elongation may mediate early dosage compensation of *btd* and *gt*. Because the MS2 and PP7 stem loop sequences in our study are inserted immediately upstream of the translation start codon and flanked by splice donor and acceptor sites from the *hb* intron, our findings speak only to the regulation of transcription initiation. However, a recent study using MS2 and PP7 reporter systems demonstrated a concerted regulation of transcription initiation as opposed to a highly variable regulation of elongation between nuclei (Liu et al. 2020). These findings suggest that elongation is not the major contributor to transcriptional regulation, though the role of elongation during early dosage compensation using these tools warrants further study.

While the primary goal of this study was to study dosage compensation, our measurements of transcription rate at endogenous loci represent a significant advance for *in vivo* studies of mechanisms of gene regulation in the *D. melanogaster* embryo. Our use of our reporter at an X-linked loci for sexing embryos also provides a straightforward control for detecting differences in the regulation of gene expression between male and female embryos.

We hope that our data, as well as the reagents generated for the purposes of our study, can help address other open questions about gene regulation in the early *D. melanogaster* embryo. The reagents generated can be used for protein colocalization studies or can be crossed mutant alleles of interest to assess the contribution of a particular molecule in the regulation of our tagged genes. Overall, we are eager to learn how the data and reagents generated in our discovery of a post-transcriptional form of early dosage compensation make possible future learnings that deepen the community’s understanding of regulating gene expression during development.

## Supporting Information

**Supplementary Figure 1: Transcription rate of *PP7-btd* and *MS2-gt* is the same in male and female embryos**

A) Distributions of fluorescent spot intensity over time (in frames, where “1” is the the first instance of transcription of the locus during nuclear cycle 14) for all female (orange) and male (dark cyan) embryos (N=9 embryos of each sex) with total spot intensity histograms on the right. B) The distribution of fluorescent spot intensities plotted over time for the dosage compensated *MS2-gt* (N=4 embryos of each sex) with total spot intensity histograms on the right. B&C) The distribution of fluorescent spot intensities plotted over time for *PP7-bnb* and *MS2-elav*, genes that are not early dosage compensated (N=3 embryos of each sex for each genotype) with total spot intensity histograms on the right.

**Supplementary Figure 2: Validation of early dosage compensation phenotype by quantitative FISH** A) A maximum intensity projection of the anterior portion of a wild type male embryo at early nc14 probed for *gt* (an early dosage compensated gene) RNA and B) a female embryo probed for *bnb* (an X-linked gene that is not early dosage compensated) RNA. Embryos were sexed by determining the number of large puncta were present within nuclei, which are dark and circular in appearance. C & D) Quantification of total probe signal for each probe in individual embryos, where the signal intensity was averaged across the expression domain with background subtraction.

**Supplemental Figure 3: Transcription rate for *PP7-btd* and *PP7-bnb* are the same in male and female embryos at nuclear cycle 13** A) The total distribution of fluorescent spot intensities and the same data plotted over the course of nuclear cycle 13 for *PP7-btd* (N=3 embryos for each sex). B) The total distribution of fluorescent spot intensities and the same data plotted over the course of nuclear cycle 13 for *PP7-bnb* (N=3 embryos for each sex).

**Supplemental Figure 4: Total distribution of spot intensities of *MS2-gt, MS2-elav*, and *PP7-bnb*** A) Total distributions of fluorescent spot intensity for all female (orange) and male (dark cyan) *MS2-gt* embryos (N=4 embryos for each sex) and the same data plotted for individual embryos by sex. We also plotted data for *MS2-elav* in B) and *PP7-bnb* in C) in the same manner (N=3 embryos for each sex per genotype).

**Supplementary Figure 5: Transcription rate of *PP7-btd* varies along the Dorsal-Ventral axis of the developing embryo** A) Distribution of fluorescent intensity of *PP7-btd* spots in individual male and female embryos. B) Variation in the width of the *PP7-btd* expression domain along the DV axis, where the dorsal (39% of the field of view) expression domain correlates with spots of weaker fluorescence intensity (indicated by the green color that matches one of the individual histograms of a male embryo in A). An embryo whose recorded *PP7-btd* expression is more ventral (52% field of view) has a wider distribution of spot fluorescent intensities. C) Distribution of fluorescent spot intensities of individual embryos matched by FOV, and therefore, DV position. D) Average particle intensities plotted over time for two male and female embryos on different positions on the DV axis.

## Acknowledgments

We thank Hernan Garcia for pioneering techniques for quantitative imaging of live embryos and for project insight; Nicholas Lammers for help with computational analysis; Gabriella Martini for generating the *MS2-gt* fly line; Barbara Meyer, Nicholas Ingolia, and Doris Bachtrog for helpful comments; Michael Stadler and members of the Eisen lab for helpful comments and project insight.

