## Supplementary figures and images for "Early dosage compensation of zygotically-expressed genes in *Drosophila melanogaster* is mediated through a post-transcriptional regulatory mechanism"

### Supplemental Figure 1

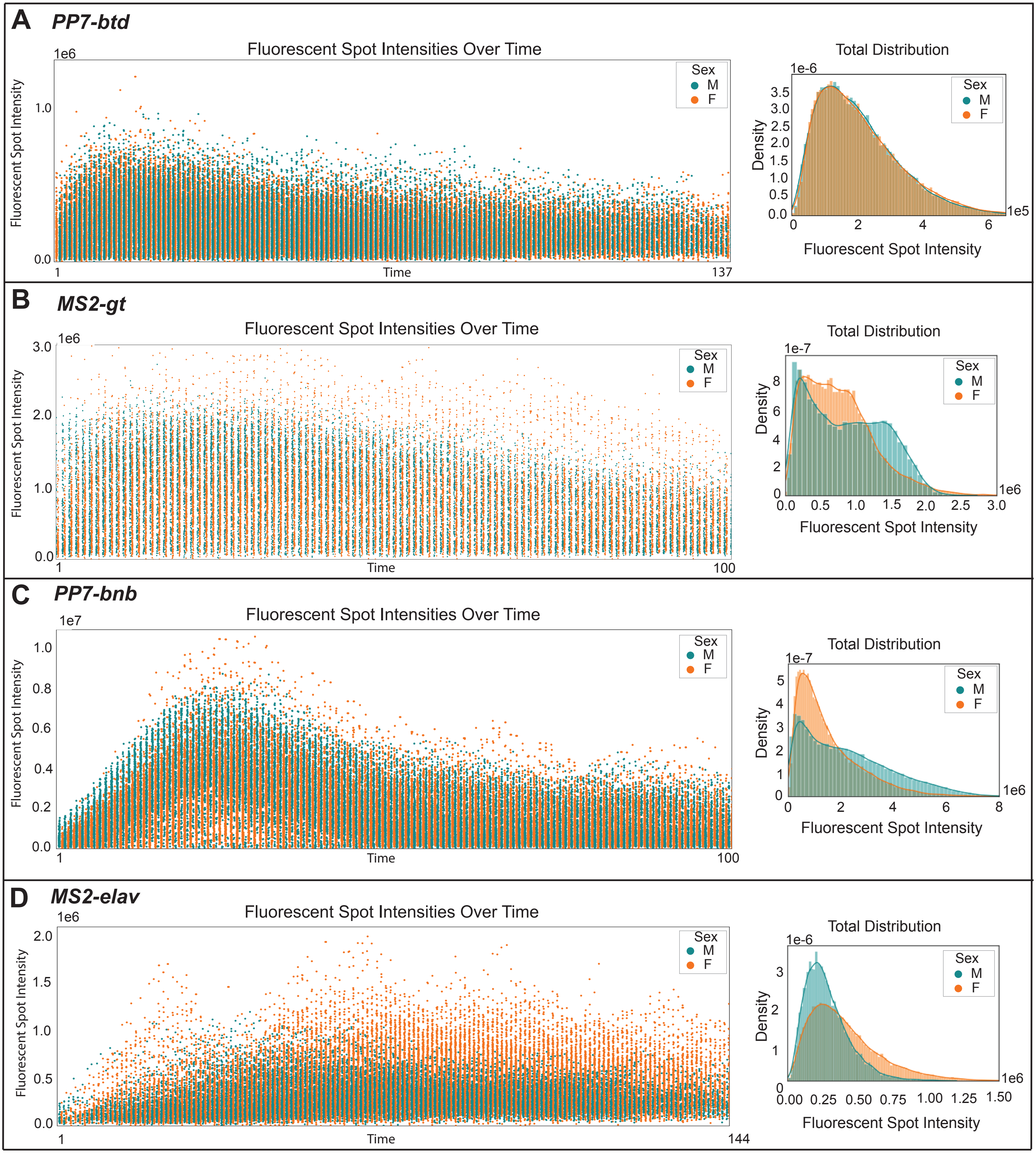

### Supplemental Figure 2

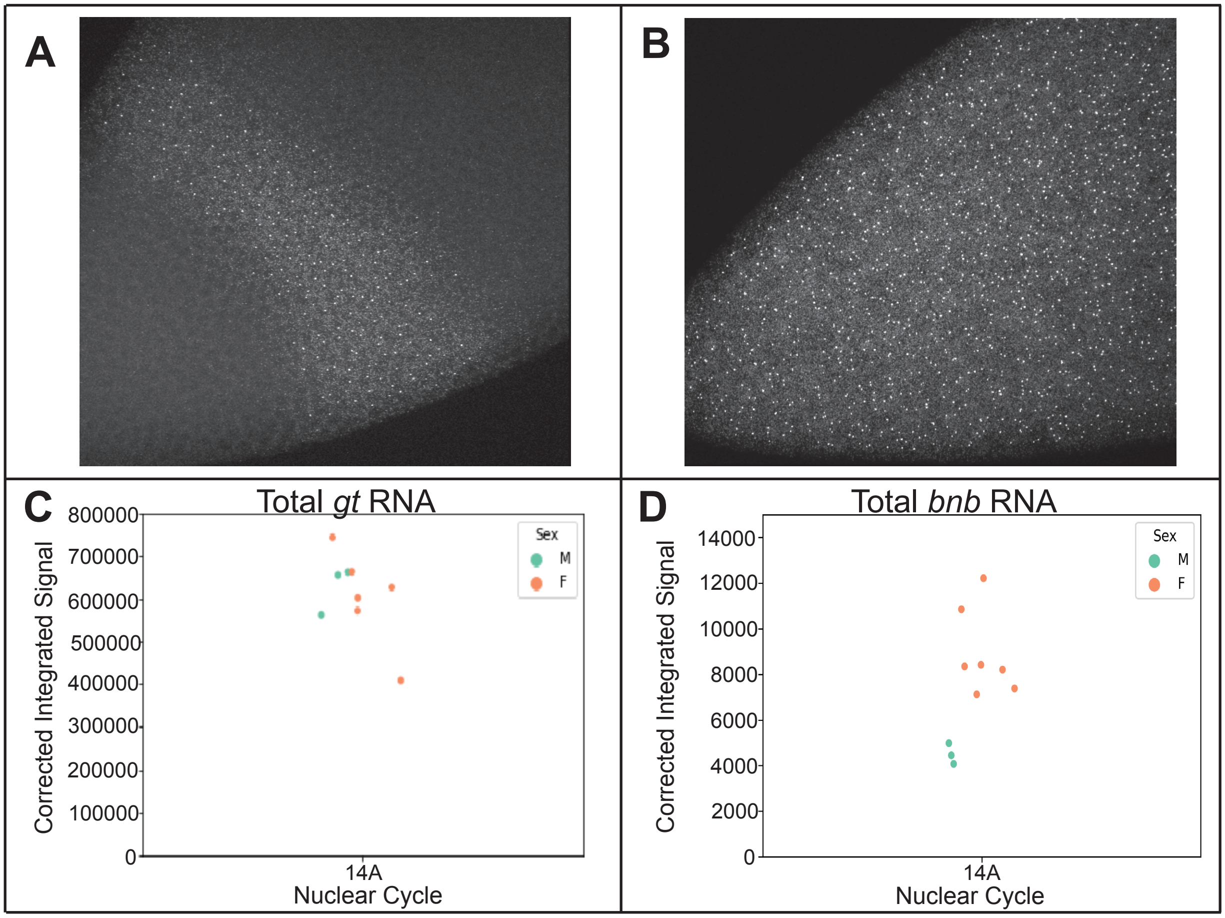

### Supplemental Figure 3

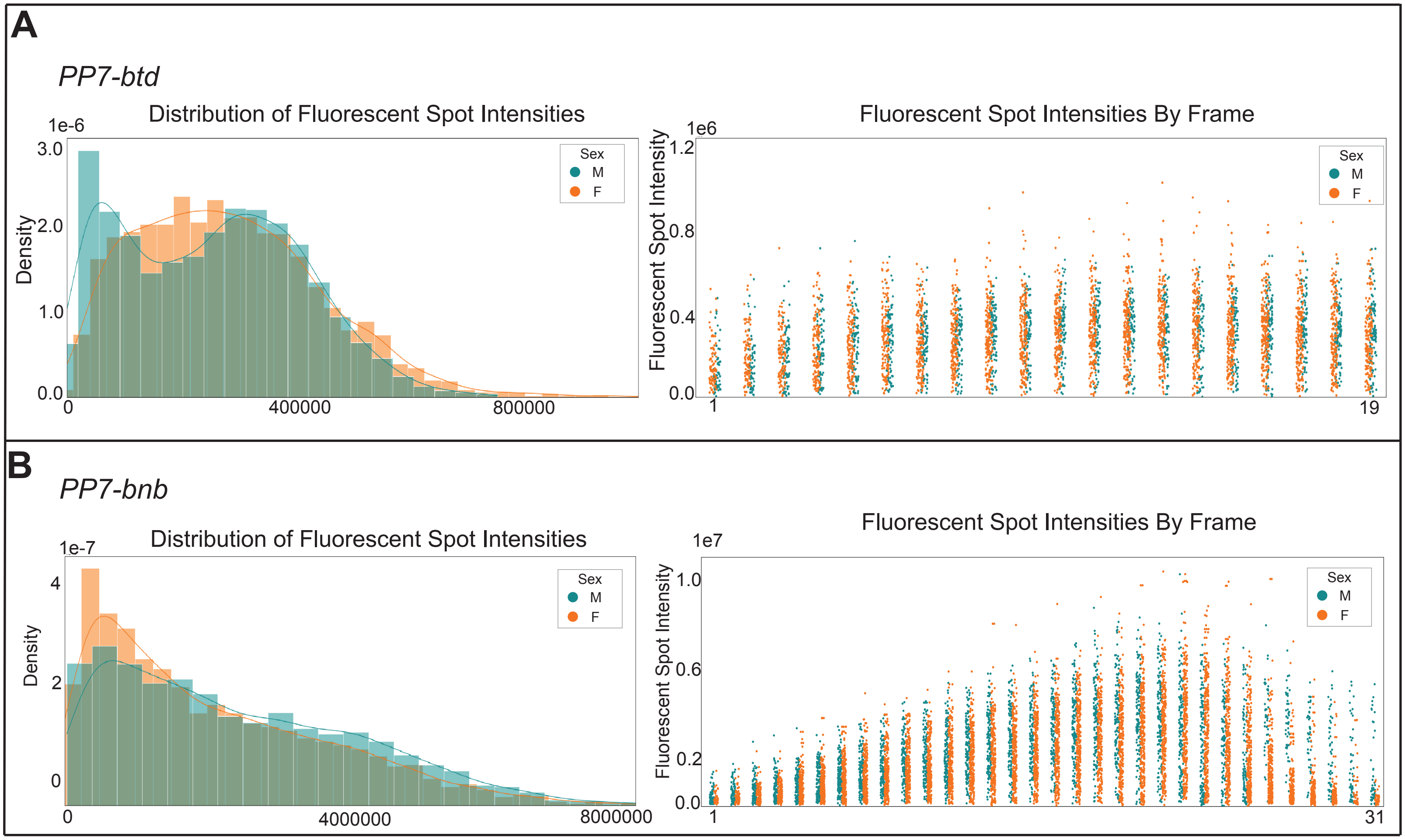

### Supplemental Figure 4

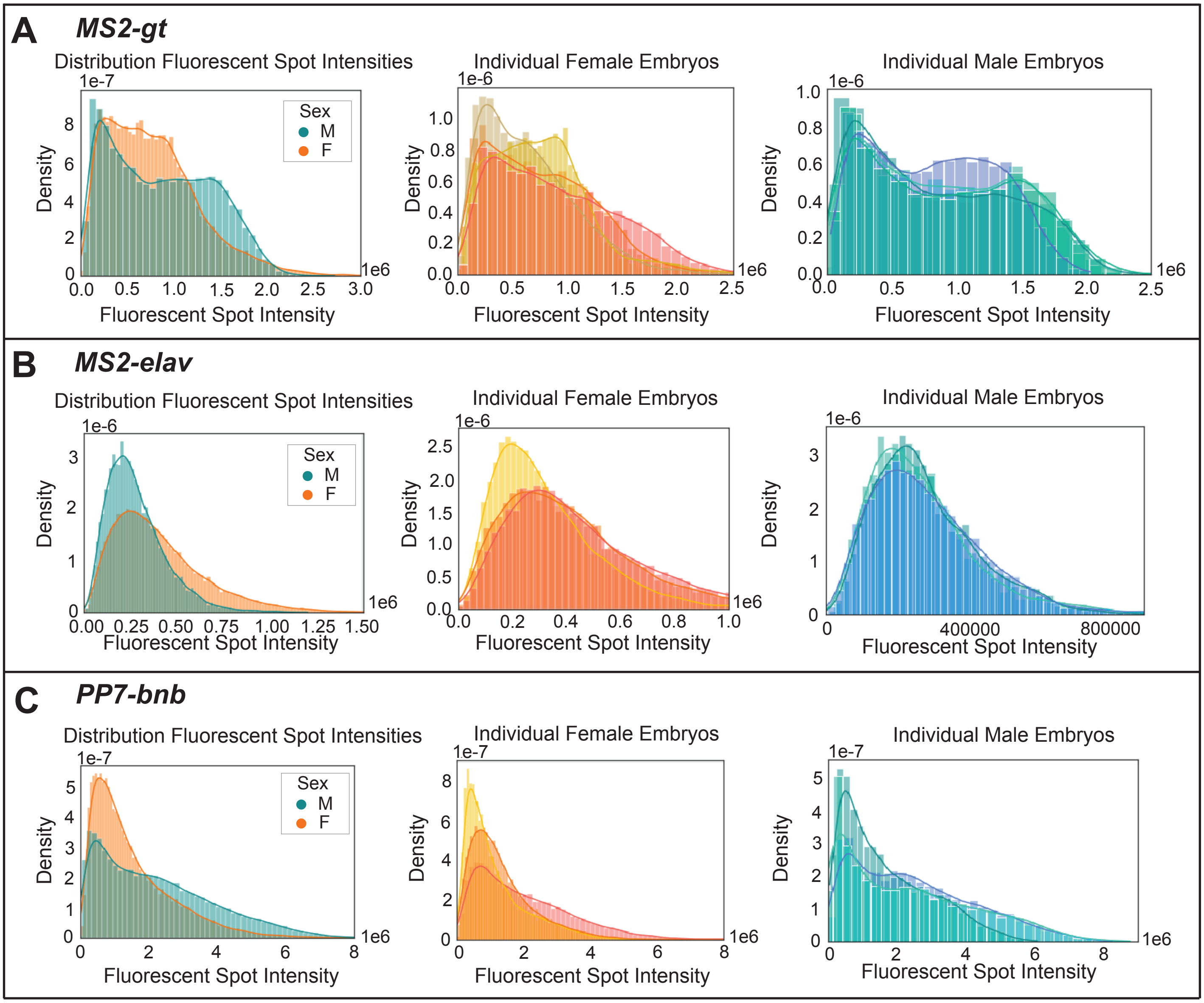

### Supplemental Figure 5

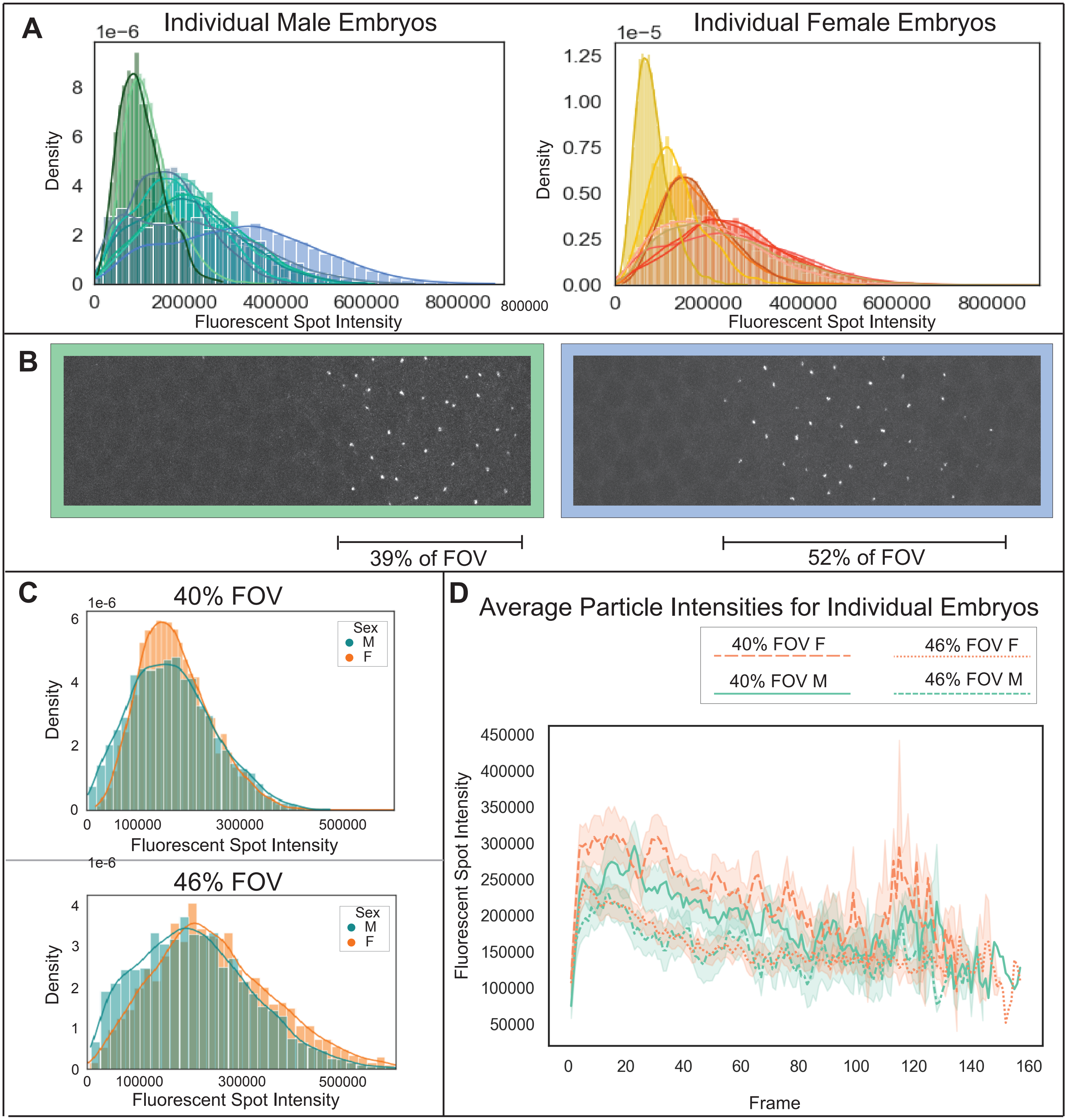
